# *miR-148a-3p* inhibits osteogenesis by targeting *Itga11* via PI3K/Akt/GSK3/β-catenin pathway

**DOI:** 10.1101/2022.10.23.513322

**Authors:** Xiaojun Chen, Kewei Rong, Xin Li, Wenqing Han, Yichuan Pang, Gang Chai

## Abstract

In previous research, miR-148a-3p deficiency was observed in bone malformation in hemifacial microsomia. Nevertheless, the mechanism of miR-148a regulating osteogenesis remains unclear. Herein, in this article, we probed into the role of miR-148a-3p in bone physiology by utilizing miR-148a knock-out (KO) mice. Compared with wild-type (WT) or heterozygotic (HE) littermates, miR-148a knock-out mice manifested lower body weight, bone dysplasia with increased bone mass. Through *in-vitro* experiments, in terms of miR-148a-3p overexpression (miRNA mimic transfection) and knockout (primary cells from WT and KO littermates), we found that miR-148a-3p can suppress osteogenesis, either in the ALP activity or bone nodules formation. Afterward, by means of proteomics, combined with RNA-sequencing and prediction databases of microRNA targets (miRDB and TargetScan), nine candidate genes targeted by miR-148a-3p were identified. Among them, only *Itga11* was regulated by mRNA degradation, while the others were modulated via post-transcriptional inhibition. Based on several online databases (GenePaint, BioGPS, STRING), *Integrin Subunit Alpha 11* (*Itga11*) was suggested to play an essential role in osteogenesis and it was confirmed as one direct target of miR-148a-3p by dual-luciferase reporter assay. Meanwhile, gene set enrichment analysis (GSEA) indicated activation of PI3K-Akt signaling pathway and WNT signaling pathway in miR-148a KO mice. The thereafter western blot confirmed that PI3K/Akt/GSK3/β-catenin signaling pathway was involved. Finally, with the help of esterase-response bone-affiliative liposome with the *Itga11* siRNA delivery system, the enhanced osteogenesis in miR-148a KO mice was partly rescued. Taken together, we demonstrated that miR-148a-3p can inhibit osteogenesis by targeting Itga11 via PI3K/Akt/GSK3/β-catenin pathway.

## Introduction

miRNAs are small endogenous non-coding RNAs composed of ∼22 nucleotides that can direct posttranscriptional modification of mRNA targets. They have been confirmed to be essential in govern stem cell fates during embryonic development^1^. They get involved in various physiological and pathological processes as well^2, 3^. In previous research, by taking advantage of the bio-samples collected from bilateral mandibles of hemifacial microsomia (OMIM: 164210) patients, miR-148a-3p was detected to be significantly decreased in the affected mandible compared with the unaffected side.

MiR-148a-3p is highly conserved in various species^4^, and has already been suggested as an important regulator in bone microenvironment. As an epigenetic regulator of bone homeostasis, miR-148a-3p was consistently upregulated in osteoporosis, and considered as a potential new plasma-based biomarker osteoporosis/ osteoporotic fracture^5^. It correlated with bone strength and metabolism as well, which had a negative correlation with total body bone mineral density^6^.

Multiple targets of miR-148a-3p have already been found based on *in vitro* experiments. Nevertheless, there was little agreement in results across studies. Moreover, complete opposite trends existed. For example, it was reported that miR-148a-3p can inhibit osteogenic differentiation via p300^7^,Wnt5a^8^, WNT1, TGFB2, IGF1^6^, NRP1^9^ or Kdm6b^10^, while it can also promote osteogenesis via SMURF1^11^. The reasons why various candidates and tendency were obtained might due to the fact that the target genes regulated by the same miRNA may vary depending on cells or tissues^2^.

In order to better understand the role of miR-148a-3p in osteogenesis, miR-148a knock-out (KO) mice were bred in this project. Meanwhile, for the first time, we probed into its regulatory mechanism in a natural *in vivo* environment.

## Materials and Methods

### Generation and genotype identification of miR-148a Knock-Out (KO) mice

miR-148a KO mice were obtained from GemPharmatech Co., Ltd. (Jiangsu, China) and housed under SPF condition in Shanghai Ninth People’s Hospital, Shanghai Jiao Tong University School of Medicine. The Ethics Committee of Shanghai Ninth People’s Hospital has approved all the following animal procedures. Heterozygous male and female mice were mating (the male-to-female ratio was 1:2) to generate miR-148a knock-out (KO), heterozygous (HE), and wild-type (WT) littermates. Ear or tail tissues were obtained for genotyping.

### Phenotype identification of miR-148a KO mice

Female mice were used here for the phenotype identification. Every mouse was weighed once a week and euthanatized at 12 weeks old when their mandibles and tibias were dissected for further examination. The length of mandibles was measured via vernier caliper, and X-ray scans were performed to assess mandibular bone mineral density. We also applied μCT to the tibias for more information about bone mass.

### Double fluorescent labeling

Calcein (8mg/kg) and Alizarin Red (20mg/kg) were administered separately by intraperitoneal injection at 10 or 11 weeks of age. These male mice were euthanatized at 12 weeks old. Mandibles were dissected and fixed in 75% ethanol. The species were then dehydrated with different concentrations of ethanol and embedded. After cutting and grinding, the sections were observed under a fluorescent microscope (OLYMPUS). The distance between the observed fluorescent bands represented the bone mineral apposition rate.

### Paraffin Section

12-week-old male mice without any treatment were sacrificed. After fixing in 4% paraformaldehyde for 48 hours, the tissues were decalcified in 10% Ethylene Diamine Tetra-acetic Acid (EDTA) for about four weeks. Then the tissues were processed by dehydrating in different concentrations of ethanol, infiltrating with xylene, and embedding with paraffin (Leica, Germany). Sections were cut at 5-μm thickness for further histochemical and immunofluorescent staining. In brief, sections were processed in a sequence of deparaffinization, hydration, antigen retrieval, permeabilization, blocking, primary antibody incubation (OCN, Servicebio, Catalog#GB11233; Runx2, Servicebio, Catalog#GB11264; Osterix, Abcam, Catalog#ab22552; ITGA11, Abcam, Catalog#ab198826), corresponding secondary antibody incubation, and nuclear staining.

### Bone Defect Animal Experiment

Eight-week-old male mice were first anesthetized by 1% pentobarbital. Then the right tibia was exposed, and muscles were dissociated in part. A pilot hole was first made with a drill point in the proximal tibia but away from the growth plate. Then the hole was enlarged with a 1 mm reamer. After irrigation, the skin was sutured. μCT was conducted on the 14^th^ day after operation to evaluate the bone healing process.

### Cell Culture

Ectomesenchymal stem cells (EMSCs) were obtained from embryos of E13.5 pregnant heterozygous mice as described in Zhao’s article^12^. In brief, the mandibular process was dissected, minced into pieces, digested with type I collagenase for two hours, followed centrifugation, resuspension and seeding. EMSCs were cultured in complete media (CM), that is, alpha-MEM supplemented with 10% Fetal Bovine Serum (FBS) and 1% penicillin/streptomycin (p./s.) (Gibco). All cultures were maintained in a 37□ incubator with 5% CO_2_.

### Cell Transfection

EMSCs were transfected with miR-148a-3p mimic reagents (RiboBio, Guangzhou, China) according to the manufacturer’s protocol. In brief, the transfection reagents were first mixed with mimic reagents. After being kept stationary for 20 minutes, the mixture was used for cell transfection for six hours. Then the mixture was replaced with CM or osteogenic media. The transfection procedure was repeated every three days if long-term culture was needed.

### Cell Proliferation Assay

Cell Counting Kit-8 (CCK8, Dojindo, China) was used to examine the role of miR-148a-3p in cell proliferation according to the manufacturer’s protocol. In brief, EMSCs were seeded on 96-well plates at a density of 3×10^3^ cells per well. At the specific time point (24h, 48h, 72h, or 96h, respectively), the standard culture medium was replaced with 100μl CM containing 10μL CCK8 reagent, and then the plates were incubated at 37□ for another two hours. Absorbances were measured at 450nm using the Absorbance Microplate Spectrophotometer (Tecan, China). Absorbance readings were normalized to the control group, and the cell proliferation curve was calculated using GraphPad Prism 8 (GraphPad Software, USA).

### Osteoblast differentiation and staining

EMSCs were maintained in CM for normal growth and osteogenic media (CM supplemented with 2 mM β-glycerophosphate, 0.05 mg/ml ascorbic acid, and 10 nM dexamethasone) for osteoblast differentiation. For miR-148a-3p mimic experiments, extra transfection procedures were done before every time the osteogenic medium was changed. Alkaline phosphatase (ALP) staining was performed on the 7^th^ day to assay the ALP activity, and Alizarin Red (AR) staining was done on the 21^st^ day to evaluate bone nodules formation. The staining protocol was as follows. Cells were first washed three times with PBS, fixed in 4% paraformaldehyde for 20 minutes at room temperature, and stained with ALP solution or 1% AR solution (Beyotime, Shanghai, China) for about 30 minutes. Cells were then washed three times with PBS again, and digital images were captured.

### Quantitative real-time polymerase chain reaction (qPCR)

For mRNA quantification, total cellular RNA was extracted from cultured cells using Multisource Total RNA Extract Reagent (Axygen, Catalog#11818KD1). The RNA purity and concentration were assessed using Nanodrop ND-1000 (Thermo Fisher). Single-stranded cDNA was reverse transcribed from 500ng total RNA using reverse transcriptase and oligo(dT) primer (Takara, Catalog#RR036). Real-time qPCR amplification was performed using the PrimeScript™ RT-PCR Kit (Takara, Catalog#RR420). *Alp, Col1a1, Runx2, Bglap, Spp1, Tnfsf11* were detected for osteogenesis. All the PCR procedures were performed in a QuantStudio 6 Flex (Applied Biosystems), and the 2^-ΔΔCT^ method was used for analysis.

Primers were as follows, *Alp* (Forward: AGAAGTTCGCTATCTGCCTTGCCT; Reverse: TGGCCAAAGGGCAATAACTAGGGA), *Col1a1* (Forward: GCTCCTCTT AGGGGCCACT; Reverse: CCACGTCTCACCATTGGGG), *Runx2* (Forward: GAC TGTGGTTACCGTCATGGC; Reverse: ACTTGGTTTTTCATAACAGCGGA); *Bglap* (Forward: TAGCAGACACCATGAGGACCATCT; Reverse: CCTGCTTGGACA TGAAGGCTTTGT), *Spp1* (Forward: AGCAAGAAACTCTTCCAAGCAA; Reverse: GTGAGATTCGTCAGATTCATCCG), *Tnfsf11* (Forward: GTACTTTCGAGCGCA GATGG; Reverse: TCCAACCATGAGCCTTCCAT).

### Western blot analysis

Total cellular protein extracts were isolated from cultured EMSCs stimulated with osteogenic media using RIPA lysis buffer (Beyotime, Catalog#P0013C) supplemented with 1% PMSF (Beyotime, Catalog#ST506). Lysates were cleared by centrifugation at 15,000*g for 10 minutes at 4°C, and the supernatant was collected. The mixture of extracted proteins and loading buffer (Beyotime, Catalog#P0015B) was denatured at 95℃ for 10 minutes and added to the prefabricated PAGE gels (GenScript, Nanjing, China). Separated proteins were then electroblotted onto polyvinylidene difluoride (PVDF) membranes (Millipore, USA). Following the transfer, membranes were blocked with 5% w/v skimmed milk in TBST for one hour and then incubated at 4□ with primary antibodies (p-PI3K, Abcam, Catalog#ab278545; p-AKT(S473), CST, Catalog#4060S; p-AKT(T308), CST, Catalog#13038S; p-GSK3b(S9), CST, Catalog#5558S; pan-PI3K, CST, Catalog#20584-1-AP; pan-AKT, CST, Catalog#4691S; pan-GSK3, Abcam, Catalog#ab185141; β-catenin, CST, Catalog#8480S; β-actin, Abcam, Catalog#ab8227) overnight. Membranes were washed thrice with TBST and then incubated with fluorescent secondary antibodies for one hour at room temperature. After washing thrice with TBST, the membrane was developed using the Odyssey infrared imaging system (Li-COR, USA).

### Omics analysis

Bones from four-week-old WT and KO littermates were collected for omics analysis. RNA sequencing was performed in order of RNA extraction (miRNeasy Mini Kit, Qiagen#217004), library construction, and sequencing with Illumina NovaSeq 6000. Proteomics was completed following protein extraction, trypsin digestion, HPLC classification, and liquid chromatography-mass spectrometry (LC-MS). Several online databases or tools were applied for further analysis, miRBD (https://mirdb.org/)^13^ and TargetScan (https://www.targetscan.org/mmu_80/)^14^ for potential targets prediction, GenePaint (https://gp3.mpg.de/)^15, 16^ for *in situ* hybridization image of E14.5 mouse embryo, BioGPS (http://biogps.org/)^17^ for candidate gene expression profiles in various tissues and cells, STRING (https://cn.string-db.org/)^18^ for the protein-protein interaction network, and jvenn (http://jvenn.toulouse.inra.fr/)^19^ for Venn diagrams.

### Dual-Luciferase Reporter Assay

Reporter plasmids containing *Itga11* 3’-UTR region with wild type or mutant miR-148a-3p binding sites (*Itga11*-WT or *Itga11*-MUT) were first constructed in pmiR-RB-REPORT™ (RiboBio). Then plasmids were co-transfected into HEK-293T cell lines with miR-148a-3p Mimic or Negative Control (NC). After culturing for 48 hours, Dual-Glo^®^ Luciferase Assay System (Promega, USA) was used to detect the luciferase activities, and firefly luciferase activities were normalized to renilla ones for the statistical analysis.

### Synthesis of the esterase-response (AspSerSer)_6_-liposome with the *Itga11* siRNA delivery system (Lipo@si-*Itga11*)

The (AspSerSer)_6_ system targeting bone-formation surfaces was first synthesis following Zhang’s experience^20^. In brief, 100mg DSPE-PEG_2000_-Maleinimide was first dissolved in 3mL DMF, then C(DSS)_6_ peptide (1.1 eq.) was added. After 12-hour reaction at room temperature, the solution was transferred to a dialysis bag (molecular weight 2000 Da) and was immersed in pure water for 24 hours. The product was collected by means of lyophilization. Afterwards, the (AspSerSer)_6_ was linked with an esterase-response liposome. In general, the PQDEA polymer was mixed with siRNA according to Prof. Shen’s report^21^. Lipids of DOPC and DOTAP were dissolved in chloroform as well as the helper like cholesterol. Then the mixture was evaporated to a film and followed by reconstructed in aqueous with added the PQDEA/DNA polyplexes dropwise. At last, the DSPE-PEG_2000_-C(DSS)_6_ was added into the system (with the ratio of 2%) and stirred for 2 hours in room temperature.

### Rescue Experiments

Six two-week-old male miR-148a KO mice were picked and randomly divided into two groups (Lipo@si-NC or Lipo@si-*Itga11*). Every three days from three to five weeks old, 100μL corresponding solution was injected via tail vein. All the mice were mercy killed at six weeks old and their tibias were dissected for further analysis, including μCT, HE staining, and immunostaining.

### Statistical analysis

As specified, all data were presented as mean ± standard deviations of three or more pooled independent experiments. Each dot in the histogram represented one independent data. Unpaired t-test was used for two groups, and analysis of variance (ANOVA) was performed for three groups or more. When ANOVA demonstrated significant differences between groups, multiple comparisons were carried out between the control group and every experimental group via Dunnett’s t-test in GraphPad Prism 8. Statistical significance was defined as * *P*<0.05, ** *P* <0.01, *** *P* <0.001 and **** *P* <0.0001.

## Results

### miR-148a Knock-Out (KO) mice manifested bone dysplasia with enhanced osteogenic potential

Since miR-148a-3p is the main product of pre-miR-148a, miR-148a KO mice (**Fig.S1a**) were utilized to investigate the function of miR-148a-3p in bone formation. Compared with heterozygotic (HE) littermates, KO mice were grossly smaller (**Fig.1a**). The body weight of 12-week-old mice was lost as the knock-out of the miR-148a allele, with an average weight of 21.92g in WT, 20.24g in HE, and 18.98g in KO (**Fig.1b**). From X-ray images, mandibular hypoplasia can be observed, both in the body length (13.83±0.27mm versus 13.36±0.34mm, *P*=0.0236) and in the ramus height (5.22±0.29mm versus 4.66±0.14mm, *P*=0.0015) (**Fig.1c**). Meanwhile, enhanced mineralization apposition rate can be observed in the KO mice (**Fig.1d**).

**Fig.1.**
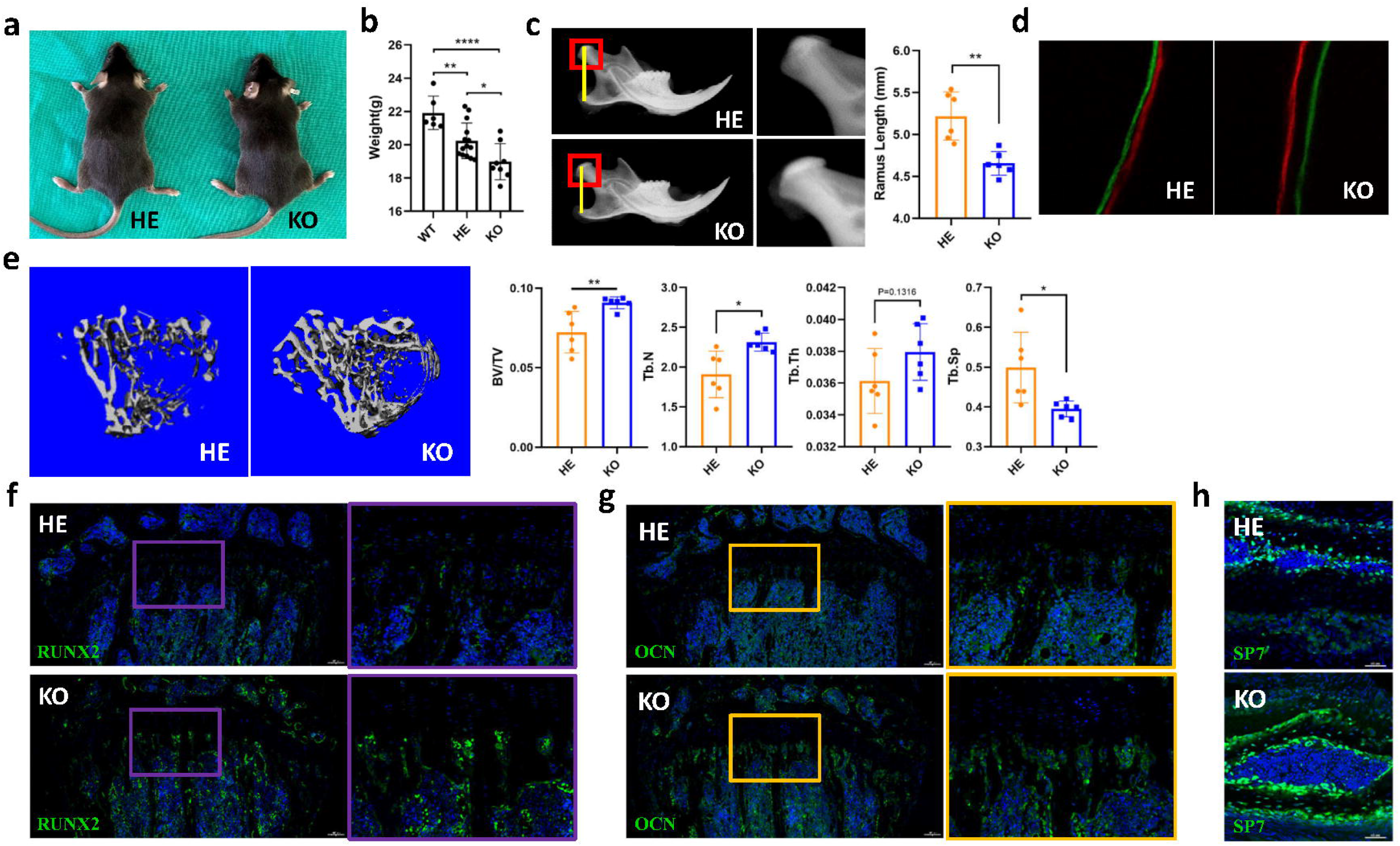
miR-148a knock-out (KO) mice manifested bone dysplasia with enhanced osteogenic potential. **(a)** Representative gross images showing knock-out (KO) mice were grossly smaller than the heterozygotic (HE) ones. **(b)** Body weights of 12-week-old mice. The weight was lost along with the knock-out of the miR-148a allele (WT, n=6; HE, n=14; KO, n=8). One-way ANOVA tests, followed by Multiple Comparisons between each column and the WT column via Dunnett’s t-test. Afterward, student t-tests were utilized between the HE and the KO group. **(c)** X-ray images (left) and length quantification (right) of mandibles from KO and HE mice (n=6 per group). Statistical analysis by student t-test. **(d)** Double fluorescent labeling of mandibles from KO and HE mice showed higher mineralization speed in KO ones. **(e)** μCT scans of tibias from knock-out (KO) and heterozygotic (HE) mice showed a remarkably higher bone mass in KO ones. Left: Representative three-dimensional reconstruction images. Right: Quantification analysis of bone parameters (n=6 per group). BV/TV, Bone Volume/Total Volume; Tb.N., Trabecular Number; Tb.Th., Trabecular Thickness; Tb.Sp., Trabecular Separation. **(f-g)** Representative immunostaining images of tibia sections (n=3 per group). **(f)** RUNX2. **(g)** OCN. **(h)** Representative Osterix (SP7)^+^ immunostaining images of cranium sections.

As the mandible largely comprises compact bone, next, we utilized long bones for further investigation. We assessed the bone mass changes through tibia μCT scans. KO mice had remarkably higher bone mass than HE ones, with increased values in bone volume/ total volume (BV/TV), trabecular number (Tb. N), trabecular thickness (Tb. Th), while decreased values in trabecular separation (Tb. Sp) (**Fig.1e**). The expression levels of Osteocalcin (OCN) and Runt-Related Transcription Factor 2 (RUNX2), both markers for osteogenesis, were significantly increased when miR-148a alleles were knocked out, especially near the growth plate (**Fig.1f**). Similar immunostaining results of Osterix (SP7) were observed in cranium (**Fig.1g**).

Bone defect animal models were also adopted, where the bone healing process was accelerated in KO mice, in comparison to WT littermates. The mean bone volume/total volume (BV/TV) increased from 0.1575 in the WT group to 0.2891 in the KO group (*P*=0.0090) (**Fig.S1b**).

### miR-148a-3p inhibited osteogenesis *in-vitro*

Primary ectomesenchymal stem cells (EMSCs) were isolated from E13.5 embryos to verify the intrinsic ability to osteogenesis *in vitro*. Compared with EMSCs from WT littermates, primary EMSCs with miR-148a KO had enhanced osteogenic potential, with more abundant and deeper ALP staining on the fourth day (**Fig.2a**).

**Fig.2.**
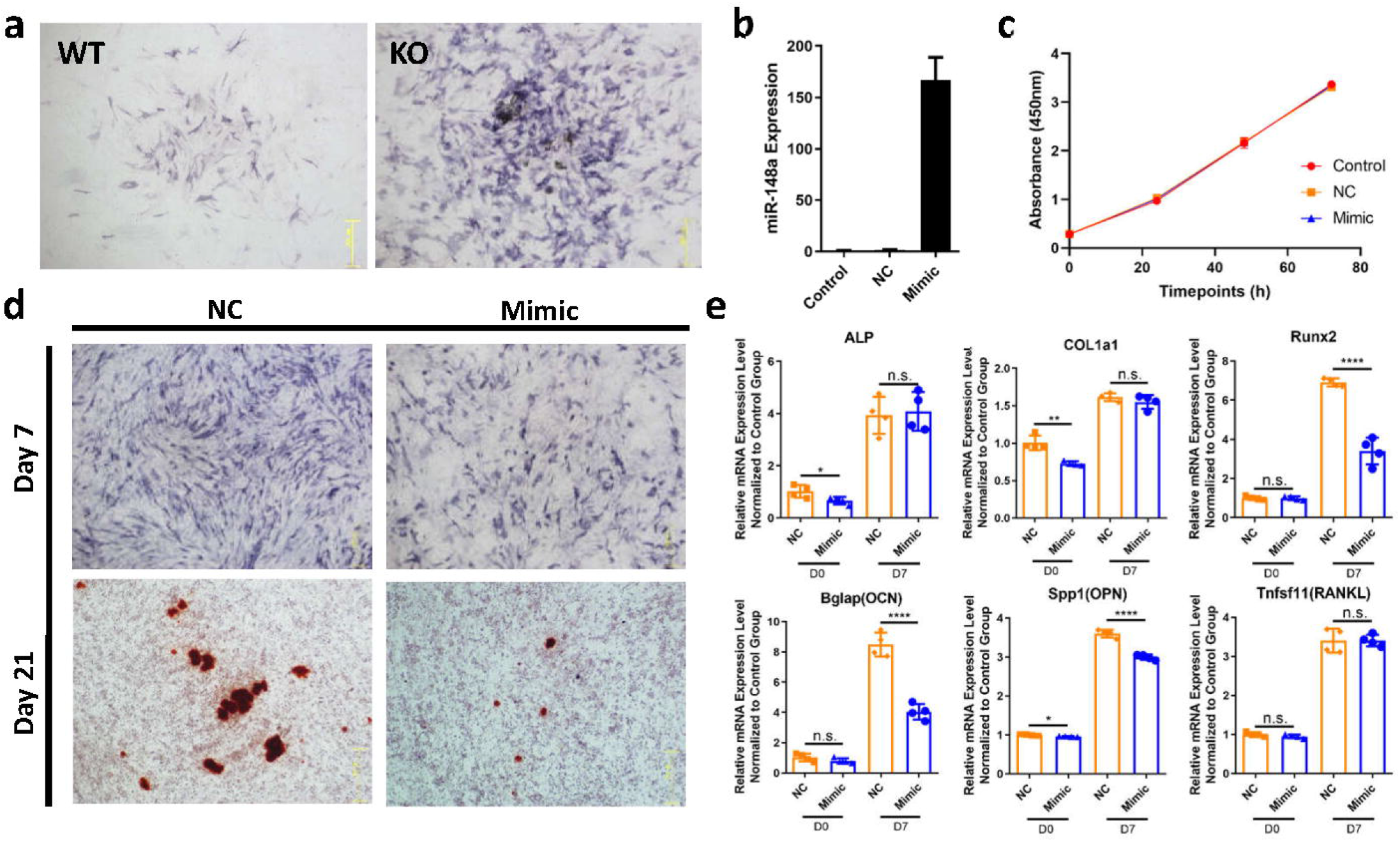
miR-148a-3p inhibited osteogenesis *in-vitro.* **(a)** Alkaline phosphatase (ALP) staining of primary ectomesenchymal stem cells (EMSCs) extracted from wild-type (WT) or knock-out (KO) embryos after 3-day osteogenic induction *in vitro*. **(b)** Validation of miR-148a-3p overexpression after miR-148a-3p mimic transfection in WT EMSCs. **(c)** Proliferation assay revealed that miR-148a-3p had no obvious influence on cell proliferation. **(d)** ALP (after 7-day osteogenic induction) and Alizarin Red (AR) staining (after 21-day osteogenic induction) of WT EMSCs with or without miR-148a-3p mimic transfection. The osteogenic potential was impaired with miR-148a-3p overexpression. **(e)** mRNA expression level after seven days’ osteogenic induction with or without miR-148a-3p overexpression. NC, Negative Control. All comparisons were performed between the NC and the Mimic group via two-sided student t-tests.

miR-148a-3p mimic was then adopted to overexpress the miR-148a-3p expression level in EMSCs extracted from WT embryos. After transfection, the miR-148a-3p expression level was significantly upregulated, over 150 folds in EMSCs (**Fig.2b**). The proliferation assay demonstrated no significant difference in cell proliferation ability among the control, NC, and Mimic groups (**Fig.2c**). Nevertheless, with miR-148a-3p overexpression, the osteogenesis was impaired, either in the ALP activity or the bone nodules formation stained by Alizarin Red solutions (**Fig.2d**). qPCR results were identical with that of staining, with lower *Alp* (*P*=0.0424), *Col1a1* (*P*=0.0016) and *Spp1* (*P*=0.0112) mRNA expression level just after transfection (without osteogenic induction), and remarkably decreased *Runx2* (*P*<0.0001), *Bglap* (*P*<0.0001) and *Spp1* (*P*<0.0001) mRNA expression level after seven days’ transfection and induction (**Fig.2e**).

### Potential targets of miR-148a-3p in regulating osteogenesis

Proteomics of bones from four-week-old WT and KO littermates were applied to study the potential mechanism of miR-148a-3p in regulating osteogenesis. By setting protein expression level ratio=1.2 (KO/WT≥1.2 or KO/WT≤0.833), 475 candidate proteins were targeted with a *P* value<0.05 (**Fig.3a**, **Supplementary Table 1**). Among them, 337 candidates were upregulated in KO mice, including ALPL (KO/WT=1.844, *P*=0.014) and SP7 (KO/WT = 2.326, *P*=0.022) (**Fig.3b**). Functional enrichment analysis of these differentially expressed proteins revealed that, compared with WT littermates, KO mice had a more powerful ossification process (**Fig.3c**), conforming to our previous *in-vivo* and *in-vitro* findings. With the help of miRDB and TargetScan database, nine candidate genes were predicted to be directly regulated by miR-148a-3p (**Fig.3d**, **Supplementary Table 2**). The protein expression levels of six candidates (CDK19, RAB34, ITGA11, PDIA3, SSR1, STT3A) were upregulated in condition of miR-148a-3p deficiency (**Fig.3e**), while the rest three (LIPA, MECP2, STXBP5) were downregulated (**Fig.S1c**).

**Fig.3.**
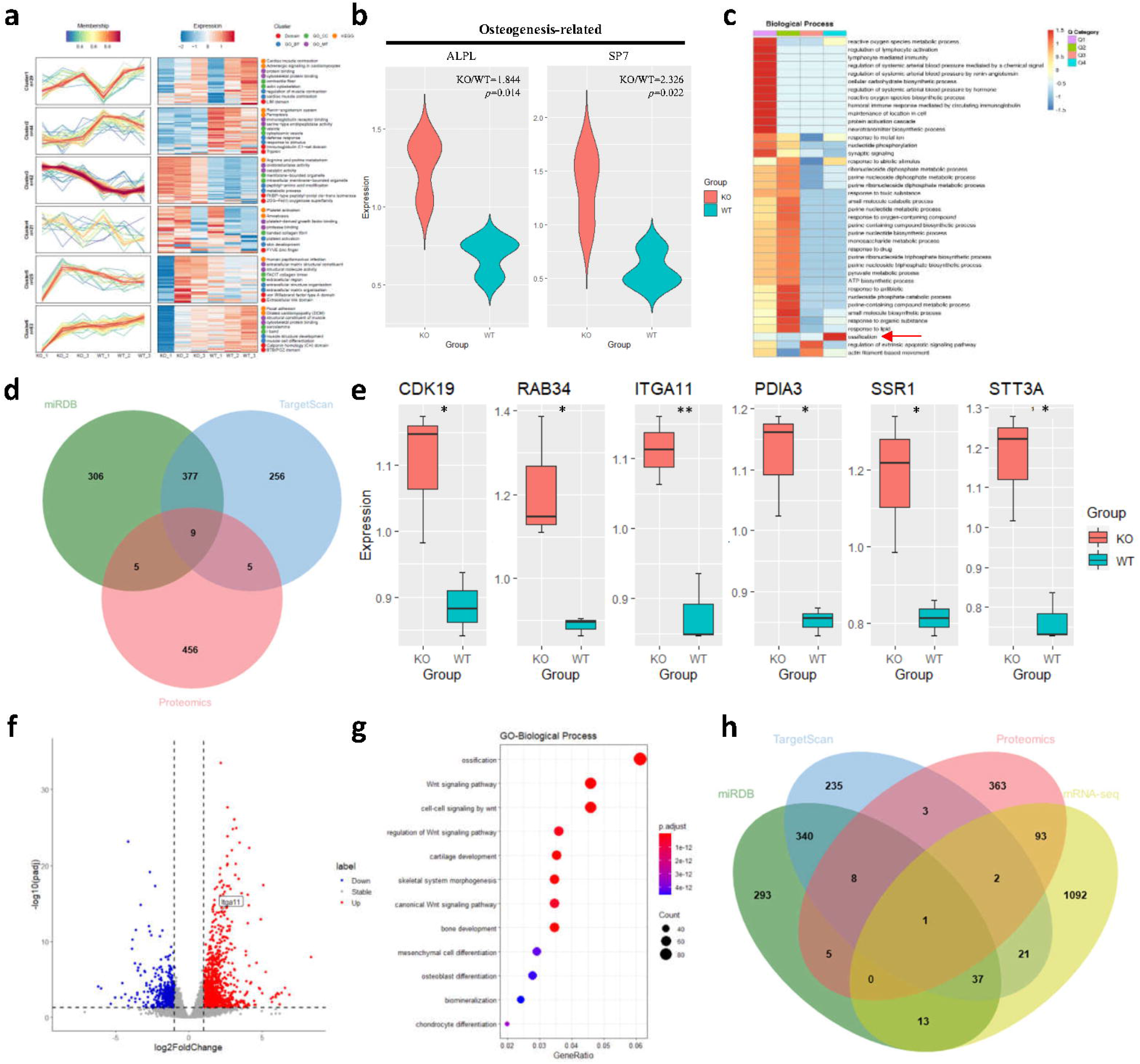
Potential targets of miR-148a-3p in regulating osteogenesis. **(a)** Proteomics results of bones from 4-week-old WT and KO littermates (n=3 per group). 475 candidates were targeted by setting protein expression level ratio=1.2 and P value<0.05. **(b)** Osteogenesis-related protein expression levels (ALPL and SP7) were upregulated in KO mice. **(c)** Functional enrichment analysis of differentially expressed proteins revealed that, compared with WT littermates, KO mice had a more powerful ossification process (Red Arrow). **(d)** Venn diagram of proteomics, miRDB, and TargetScan results. Nine candidates that miR-148a-3p may directly regulate were found. **(e)** The protein expression levels of six candidates (CDK19, RAB34, ITGA11, PDIA3, SSR1, STT3A) were upregulated in condition of miR-148a-3p deficiency. **(f)** mRNA-sequencing results of bones from WT and KO littermates (n=3 per group) when Q<0.05 and FC2 were set as filter parameters. **(g)** GO enrichment found that “Ossification” was greatly enhanced in KO mice. **(h)** Venn diagram of proteomics, mRNA-sequencing, miRDB, and TargetScan results, indicating miR-148a-3p can modulate ITGA11 protein expression level by degrading its mRNA.

Traditionally, miRNAs suppress the post-transcriptional expression of their target genes via two primary mechanisms, either through mRNA degradation or not^2^. Thus, in order to clarify the exact mechanism of how miR-148a regulate these candidates, mRNA-sequencing was accomplished to find out the gene mRNA expression differences between KO and WT littermates. When Q<0.05 and FC2 (KO/WT≥2 or ≤0.5) were set as filter parameters, 1259 genes were found in all (**Fig.3f**, **Supplementary Table 3**), among which 879 genes were upregulated. Once again, GO enrichment found that “Ossification” was greatly enhanced in KO mice (**Fig.3g**). The mRNA-seq results were then combined into the Venn diagram, only one candidate, *Itga11*, remained (**Fig.3h**, **Supplementary Table 4**), indicating miR-148a-3p can completely complement with Itga11 3’-UTR and modulate its protein expression level by mRNA cleavage. In comparison, the rest candidates were probably regulated via post-translational and translational repression.

### miR-148a inhibits osteogenesis by targeting Itga11

Based on several online databases, Itga11 was suggested to play an essential role in osteogenesis. The *in situ* hybridization image of E14.5 mouse embryo (obtained from GenePaint, https://gp3.mpg.de/) demonstrated high expression level of Itga11 in Meckel’s cartilage, ribs and spine during embryonic period (**Fig.4a**). The expression pattern of Itga11 (obtained from BioGPS, http://biogps.org/) revealed that Itga11 was highly expressed in osteoblasts, over 30 times more than in other tissues or cells (**Fig.4b**). In addition, the protein-protein interaction network (obtained from STRING, https://cn.string-db.org/) showed interaction between Itga11 and Type □ Collagen (Col1a1 and Col1a2), a specific marker in osteogenesis (**Fig.4c**). What’s more, Itga11 has already been reported as a regulator in bone homeostasis, as a receptor for either Collagen XIII^22^ or osteolectin^23^.

**Fig.4.**
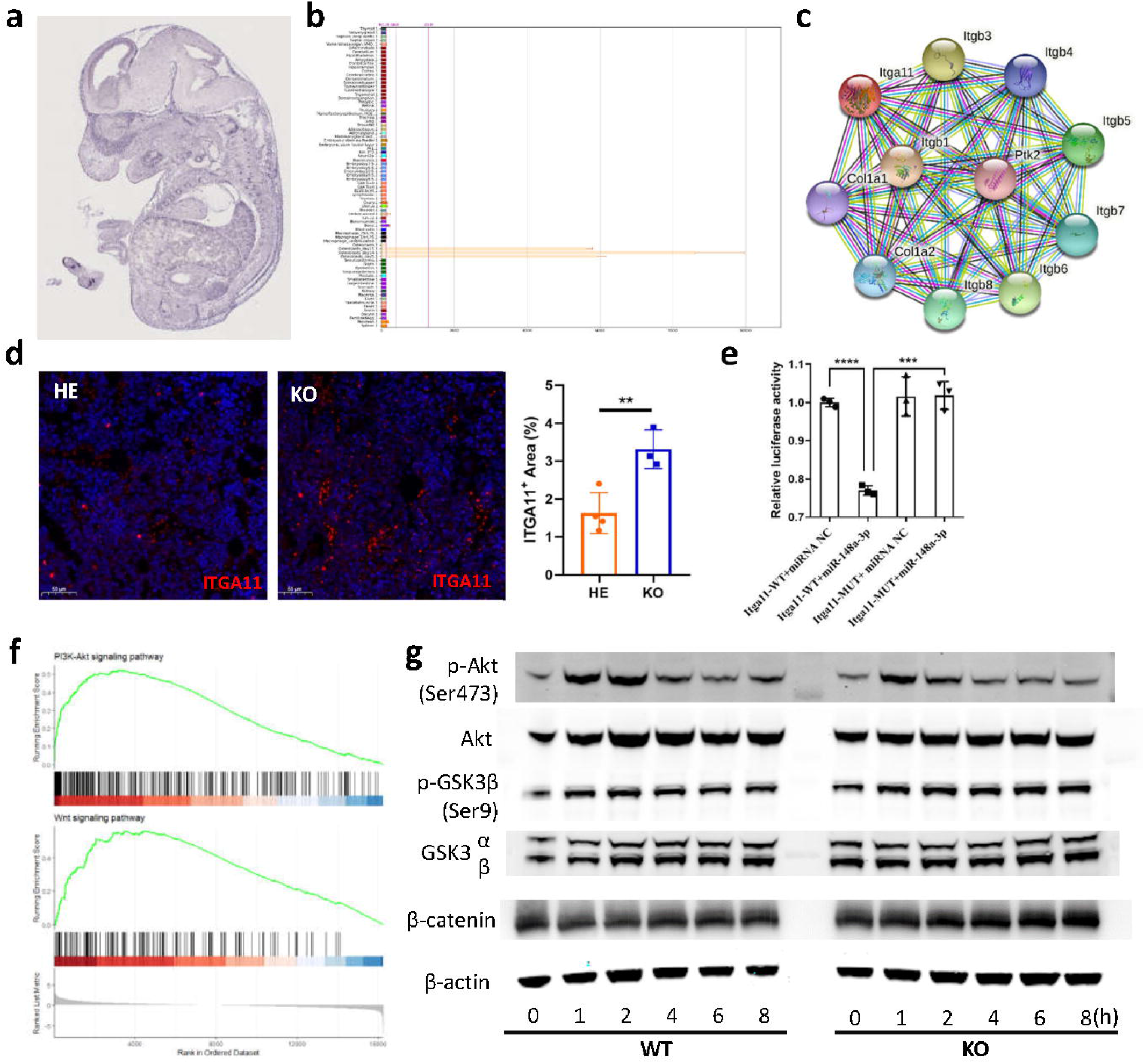
miR-148a inhibit osteogenesis by targeting *Itga11* via PI3K/Akt/GSK3/β-catenin signaling pathway. **(a)** The *in situ* hybridization image of Itga11 in E14.5 mouse embryo obtained from GenePaint. **(b)** Itga11 gene expression profile of Itga11 obtained from BioGPS, with a rather high expression in osteoblasts. **(c)** The protein-protein interaction network obtained from STRING showed interaction between Itga11 and Type □ Collagen, a specific marker in osteogenesis. **(d)** Immunostaining results verified the higher ITGA11 expression in KO littermates. **(e)** Dual-luciferase reporter assay confirmed that miR-148a-3p could bind to the 3’UTR of Itga11 and regulate its expression. **(f)** Gene set enrichment analysis (GSEA) of mRNA-sequencing results indicated activation of PI3K/Akt and Wnt signaling pathway. **(g)** PI3K/Akt/GSK3/β-catenin signaling pathway was promoted in condition of miR-148a knock-out.

Next, immunostaining results verified the higher expression in KO littermates, in comparison to HE ones (**Fig.4d**). Dual-luciferase reporter assay verified the direct combination of miR-148a-3p and *Itga11* 3’-Untranslated Regions (3’-UTR). The luciferase activity was reduced by 23% when supplemented with miR-148a-3p and *Itga11*-WT (*P*=0.0004), while almost no difference with miRNA NC or *Itga11*-MUT (**Fig.4e**).

### miR-148a inhibits osteogenesis via PI3K/Akt/GSK3/β-catenin signaling pathway

Gene set enrichment analysis (GSEA) of the abovementioned mRNA-sequencing results indicated activation of some key metabolic pathways, including PI3K-Akt signaling pathway and Wnt signaling pathway (**Fig.4f**). Similar results were observed in proteomics. Key proteins, such as AKT and β-catenin, were remarkably upregulated in miR-148a KO samples (**Fig.S1d**).

The molecular mechanism was further studied through western blot. With the stimulation of osteogenic media, the phosphorylation proportion of AKT (p-AKT/ pan-AKT) was obviously increased and reached climax at one hour at a level of about 0.096, then decayed with time. In the absence of miR-148a, the climax time point was not changed but peak value raised to 0.105. In the WT group, the phosphorylation proportion of GSK3β (p-GSK3β/pan-GSK3β), the inactive form of GSK3β, reached peak ratio of 0.801 at 2-hour stimulation. Then the values came down over time. While in the KO group, it remained at a relative high level till the eighth hour (0.907 at the second hour and 0.659 at the eighth hour). Meanwhile, obvious increasement can be observed in β-catenin protein expression level, a terminal protein of the classic WNT signaling pathway^24^ (**Fig.4g**).

### *Itga11* silencing partly rescued the excessive osteogenesis in miR-148a KO mice

Although *Itga11* was highly expressed in osteoblasts, it was more or less expressed by other cells or tissues as well. Herein, we introduced a bone-affiliative (AspSerSer)_6_-liposome-based siRNA delivery system (Lipo@siRNA) that can specifically transport siRNA to skeletal tissues by intracellular esterase-response (**Fig.5a, Fig.S2**) and tried rescuing the excessive osteogenesis in miR-148a KO mice via Lipo@si-*Itga11* tail vein injection (**Fig.5b**).

**Fig.5.**
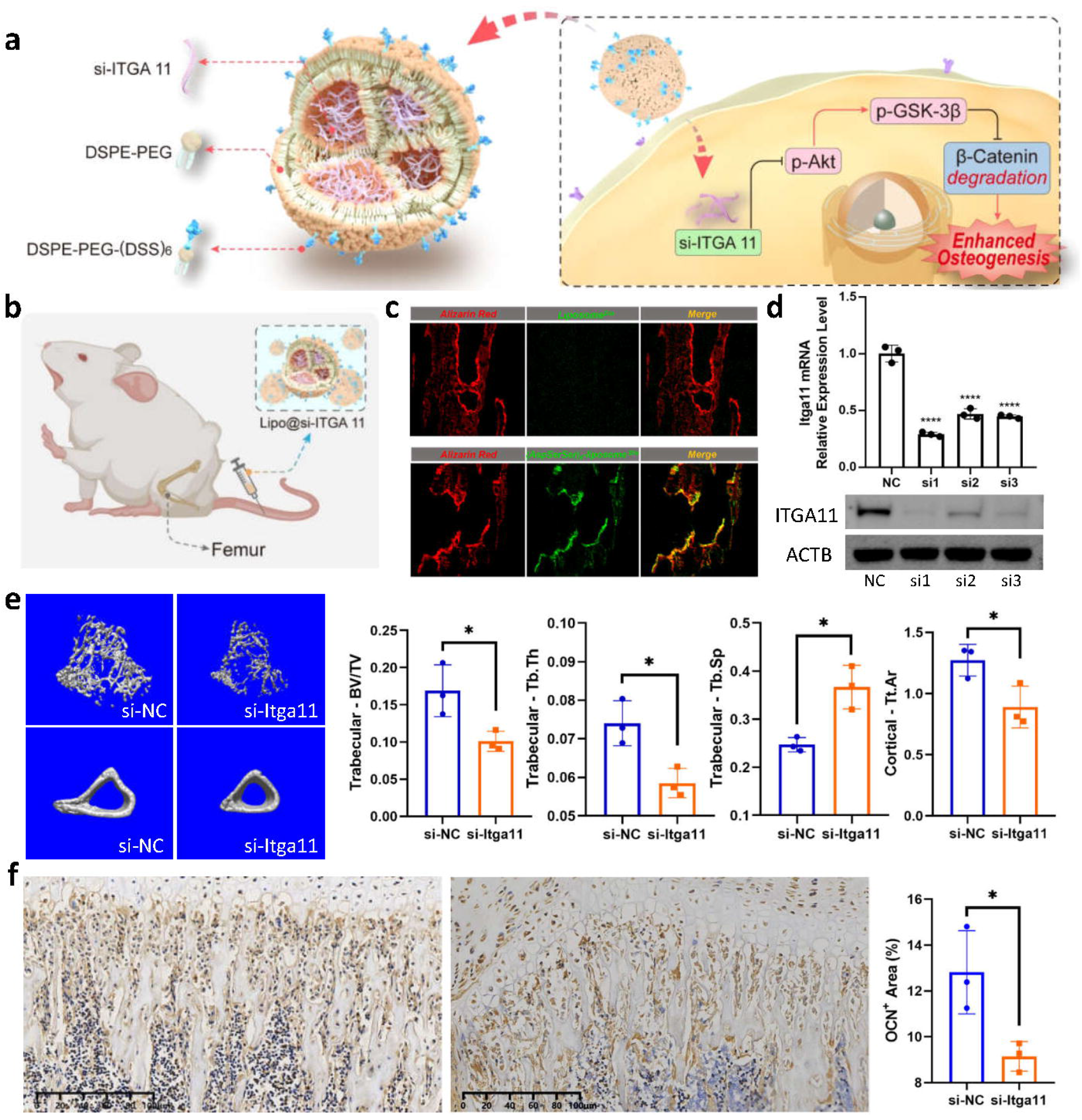
ITGA11 silencing partly rescued the excessive osteogenesis in miR-148a KO mice. **(a)** The schematic diagram of the esterase-response (AspSerSer)_6_-liposome-based siRNA delivery system (Lipo@si-ITGA11). **(b)** The sketch map of the rescue experiments via Lipo@si-ITGA11 tail vein injection. **(c)** The validation of the bone-targeting design by means of undecalcificated bone slicing. **(d)** The silencing efficiency of three different siRNA sequences. Upper: mRNA expression level; Below: protein expression level. **(e-f)** Silencing overexpressed ITGA11 by Lipo@si-ITGA11 in miR-148a KO mice can partly rescue the excessive osteogenesis (n=3 in each group). **(e)** μ-CT images and quantitative parameters. BV/TV, trabecular bone volume/ total volume. Tb.Th, trabecular thickness. Tb.Sp, trabecular separation. Tt.Ar, cortical total area. **(f)** The representative images and quantitative analysis of osteocalcin (OCN) immunostaining.

By means of undecalcificated bone slicing, we found that the green fluorescence signals of (AspSerSer)_6_-liposome (labeled by Dio) were mainly distributed along the trabecula (marked by Alizarin Red) in bones after intravenous injection, conforming to its bone targeting design. While little signals can be observed in the traditional liposome group (**Fig.5c**).

Afterwards, three different siRNA sequences were composited to downregulate the *Itga11* expression level. All the sequences were effective. Compared with si-NC group, the silencing efficiency was 71.33% in si1 group, 53.33% in si2 group and 55.67% in si3 group, separately (All P<0.0001). Similar results were obtained when it regards to protein expression level (**Fig.5d**). Therefore, si1 was used in the Lipo@*si-Itga11* system and our following research.

By means of μ-CT and immunostaining, *in-vivo* data revealed that silencing overexpressed ITGA11 in miR-148a KO mice can partly rescue the abnormal osteogenesis. Compared with the Lipo@si-NC group, the ones in Lipo@si-*Itga11* group possess parameters of decreased osteogenic ability, such as declined trabecular bone volume/ total volume (BV/TV) (P=0.0346), thinner trabecular thickness (Tb.Th) (P=0.00182), increased trabecular separation (Tb.Sp) (P=0.0122) and reduced cortical total area (Tt.Ar) (P=0.0365) (**Fig.5e**). Besides, the immunostaining results of osteocalcin (OCN), whose average positive stained area percentage dropped from 12.810% to 9.147% (P=0.0300) (**Fig.5f**), also indicated that the excessive osteogenesis in miR-148a KO mice was partly resolved via ITGA11 silencing.

## Discussion

In this article, we utilized miR-148a KO mice to study the role of miR-148a-3p in bone physiology. By taking advantage of miR-148a KO mice, we assessed the role of miR-148a in bone physiology *in vivo* for the first time. miR-148a KO mice manifested bone dysplasia with increased bone mass. Meanwhile, proteomics and RNA-sequencing of bone samples from littermates were performed to look for candidate targets in a natural *in vivo* environment. Itga11 was verified to be one direct target of miR-148a-3p in inhibiting osteogenesis via PI3K/Akt/GSK3/β-catenin signaling pathway. Lastly, by taking advantage of a bone-targeting siRNA delivery system, the excessive osteogenesis was partly rescued by silencing ITGA11, which proved that ITGA11 was one direct target of miR-148a in regulating osteogenesis and provide new therapeutic options.

Multiple targets of miR-148a-3p that regulate bone physiology have already been reported. However, most of them were identified based on cell lines through *in vitro* experiments. Taking osteogenesis regulation for example, various target genes existed, like p300^7^,Wnt5a^8^, WNT1, TGFB2, IGF1^6^, NRP1^9^, Kdm6b^10^, and SMURF1^11^. Due to the nature that the target genes regulated by the same miRNA may vary depending on cells or tissues^2^, looking for targets based on *in vivo* environment might be a better choice. Utilizing bone samples from four-week-old littermates, *Itga11* was found as the main effector of miR-148a in regulating osteogenesis. As reported, *Itga11*, a marker of mesenchymal stem cells^25^, has an essential role during embryogenesis, especially in mesenchymal cells around the cartilage anlage in the developing skeleton^26, 27^, and takes part in the regulation of bone homeostasis^28, 29^. Itga11 has also been reported to be involved in PI3K/Akt and Wnt signaling pathway, supporting our findings from another view. ITGA11 is one of the core genes that could influence PI3K/Akt signaling pathway once they were activated^30^, its expression can affect tumor progression by modulating the PI3K/Akt pathway^31^. Besides, Wnt pathway can be activated by osteolectin/α11β1 signaling^29^.

Nevertheless, we failed to unveil the mechanism of bone dysplasia, especially in aspect of shortened length. Osteoclastogenesis may get involved. Besides, according to recent publication, it might be related to abnormal endothelial proteolytic activities, which has been proved to have a vital role in bone elongation instead of mature osteoclasts^32^. ALDH18A1, which was defined as an important candidate in pathological angiogenesis through single-cell RNA sequencing^33^, was significantly increased in KO mice (KO/WT=1.471, *P*=0.0119). Literatures have revealed the role of miR-148a in regulating angiogenesis^34-42^ and osteoclastogenesis^43, 44^. Further related researches should be carried out. What’s more, crosstalk between genes and unconventional functions of miRNA^45^ should be taken into consideration as well.

In summary, our findings showed that miR-148a-3p inhibited osteogenesis by targeting Itga11 via PI3K/Akt/GSK3/β-catenin pathway, which might partly be the mechanism of how miR-148a deficiency caused bone dysplasia.

## Acknowledgements

This work was supported by National Natural Science Foundation of China (81741012); Shanghai Municipal Key Clinical Specialty-shslczdzk00901 (ZWJCB18); Shanghai Sailing Program (23YF1421700). Thank PTM BIOLABS for their great help in proteomics analysis.

## Author contributions

XJ Chen, YC Pang and G Chai designed research; XJ Chen, RK Wei, X Li, and WQ Han performed research; YC Pang constructed the delivery system; XJ Chen and YC Pang analyzed the data; XJ Chen drafted the paper; YC Pang and G Chai revised the paper.

## Supplementary Figure Legends

**Fig.S1** (a) miRNA-sequencing of bone samples from littermates confirmed little miR-148a-3p/5p expression in miR-148a KO mice. **(b)** Bone defect animal models indicated an accelerated bone healing process in KO mice in comparison to WT ones. Left: Representative middle-cross section of μCT scans. Right: Quantification of the bone defect site (n=6 per group). **(c)** The protein expression levels of three candidates (LIPA, MECP2, STXBP5) were downregulated in condition of miR-148a-3p deficiency. **(d)** Protein expression pattern in PI3K/Akt and Wnt signaling pathway obtained from proteomics. Key proteins, such as AKT and β-catenin, were remarkably upregulated in miR-148a KO samples. Red box: Upregulated proteins; Green box: Downregulated proteins.

**Fig.S2** (a) The blueprint of the esterase-response (AspSerSer)6-liposome-based siRNA delivery system (Lipo@siRNA). **(b-c)** The compound of DSPE-PEG_2000_-C(AspSerSer)_6_. **(b)** The chemical equation. **(c)** The Nuclear Magnetic Resonance. **(d-e)** The constitution of the esterase-response system. **(d)** The chemical equation. **(e)** The Nuclear Magnetic Resonance.

## Supplementary Table

**Supplementary Table 1. Proteomics results of bones from 4-week-old knock-out (KO) and wild-type (WT) littermates.**

**Supplementary Table 2. Raw data of Venn diagram composed of proteomics, miRDB and TargetScan results.**

**Supplementary Table 3. mRNA-sequencing results of bones from 4-week-old knock-out (KO) and wild-type (WT) littermates.**

**Supplementary Table 4. Raw data of Venn diagram composed of proteomics, mRNA-sequencing, miRDB and TargetScan results.**

