## Supplementary figures and images for "*miR-148a-3p* inhibits osteogenesis by targeting *Itga11* via PI3K/Akt/GSK3/β-catenin pathway"

### Fig.S1

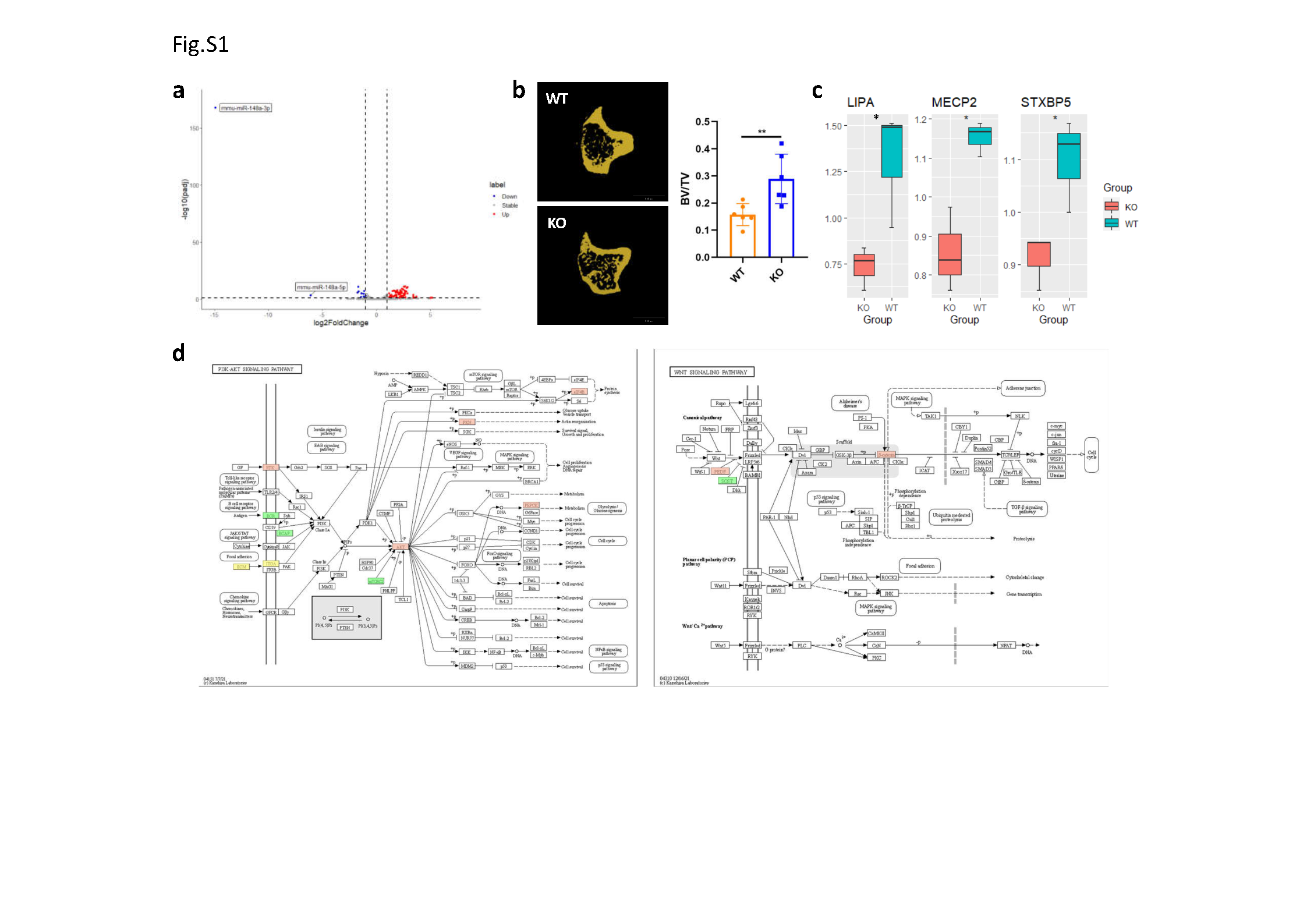

### Fig.S2

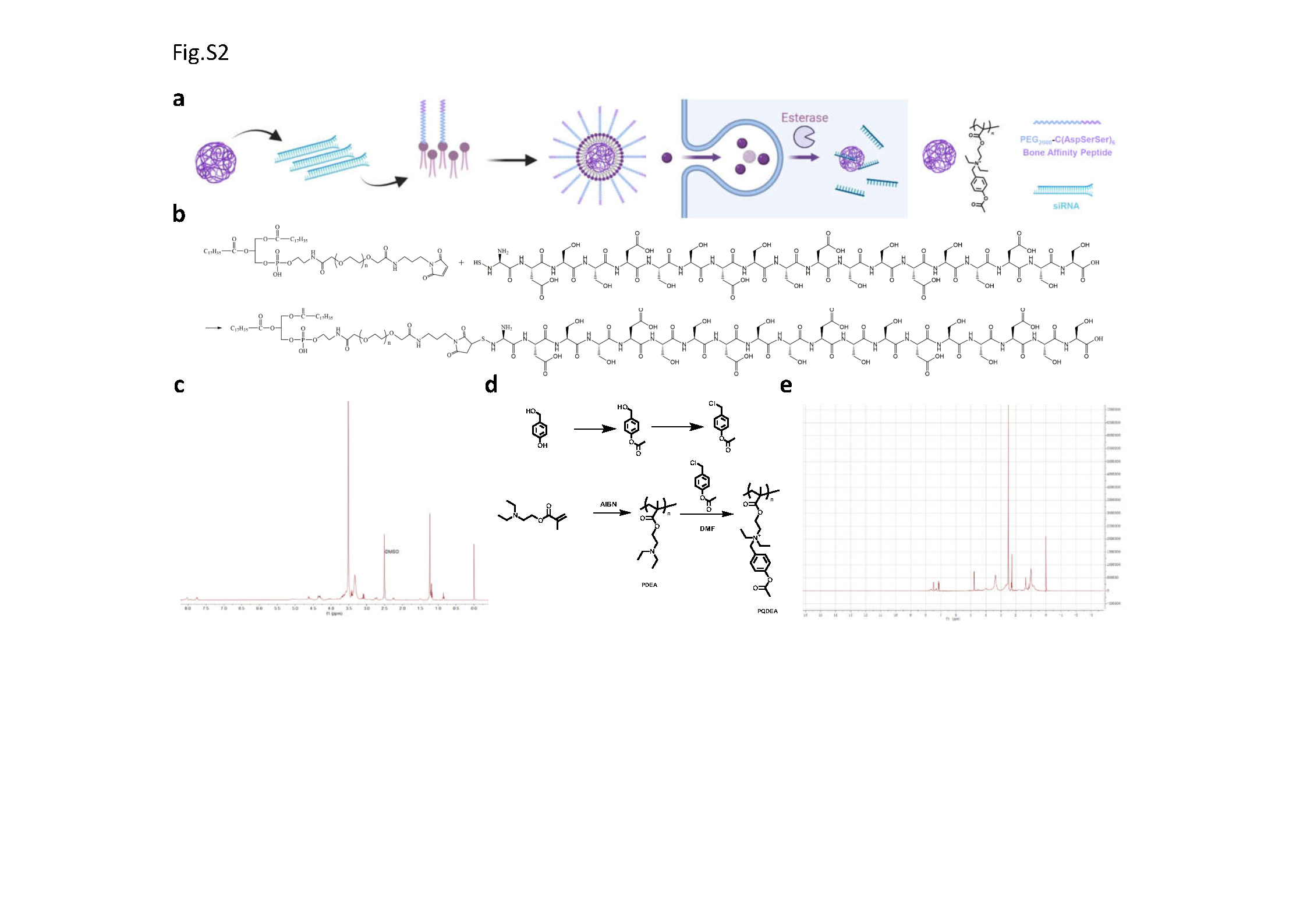
